# Phumdis-A Promising Alternative Energy Resource

**DOI:** 10.1101/2021.04.12.439410

**Authors:** Ayesha Siddiqua, Sweekrity Kanodia, Jincy Jacob, Darwilin Khumanthem, Keke Thakhell, C.K Chandra Babu, Saisha Vinjamuri

**Author notes:** Corresponding author : Saisha Vinjamuri, e-mail id.

## Abstract

The inadequate sources of energy coupled with the increasing demands of power have necessitated the search for novel renewable energy resources. ‘Phumdis’ are one such promising alternative. Phumdis are floating mats of heterogeneous mass of vegetation, soil and organic matter found in Loktak Lake in Manipur, North Eastern India. This paper delineates the use of Phumdis as an alternative energy source. Phumdis from Loktak Lake, Manipur were processed and analyzed for their biofuel capabilities. The Results indicate that the phumdis have high calorific content, cloudpoint and flash point indicating that they are at par with other fuels.

## 1. INTRODUCTION

The fast depleting petroleum reserves and environmental concerns have led to the search for alternative environmentally friendly renewable sources of energy. (Georgogianni et al., 2008; Meher et al., 2006; Narasimharao et al., 2007). Though biodiesel from algae is considered as the most suitable alternative (McKendry,2002; Reijnders, 2006), the processes for its cultivation and harvesting the biodiesel is still in its infancy and is currently not sufficient to meet the increasing demands of energy(Torzillo, et al 2003.) This has led to a search for alternative sustainable resources. Here we report the use of phumdis as a promising alternative.

“Phumdis” are floating islands of vegetation, extensively found in Lotak Lake, Manipur, North East India. They are a heterogeneous mass of soil, vegetation and organic matter at various stages of decomposition (Trisal&Manihar, 2004) the core of which is composed of black colored detritus. Of late there has been an increase in proliferation of Phumdis resulting in impeded water circulation and eutrophication of the lake. This has necessitated their removal from time to time. The use of phumdis for biofuel production promises to be win – win situation by helping in maintaining the lake ecosystem as well as generating biofuel.

## 2. MATERIAL AND METHODS

### 2.1 Collection of Plant Material

Phumdis (10kg) were collected from Loktak Lake, Manipur, washed to remove debris, separated into different parts viz roots & stems and air dried. The dried samples were finely powdered and stored in a cool dry place till further use.

### 2.2 Extraction, transesterification and FAME analysis

Extraction and estimation of total lipid content of the three powdered samples viz roots and stems, was carried out as per the procedures given by Bligh and Dyer (1959). To the extracted samples, sodium hydroxide (NaOH) : methanol (1:400) mixture was added e in the ratio 6:1 to initiate the transesterification process. This mixture was stirred for an hour and then allowed to settle for 2 days so as to obtain 2 separate layers; the supernatant layer (biodiesel) and sediment layers (methanol). The biodiesel was separated carefully from the sediment layer by a flask separator and methanol was removed using a rotary evaporator at 50°C. The biodiesel obtained was stored and the amount of fatty acid methyl esters (FAMEs) were determined using a gas chromatography mass spectrometer (GCMS)(Agilent systems). The conditions used include hydrogen as the carrier gas with a flow rate of 40ml/minutes. The Gas chromatography column was set at 220°C as its maximum temperature. The initial temperature of the column was at 80°C. Thermal gradient was 220°c at a rate of 13°C/ minutes. The post temperature reached was 50°C at 2minutes.

### 2.3 Analysis of Flash Point, Gross Calorific Value, and Cloud Point

#### 2.3.1 Flash point

It is the lowest temperature of the sample, corrected to a barometric pressure of 101.3 kPa, at which application of a test flame causes the vapour of the sample to ignite (Indian Standard Methods of Test for Petroleum and its Products (P20)). The flash point determines the flammability of a material. Abel’s closed cup was used for this experiment. The sample, suitably cooled, was placed in the cup of the Abel apparatus and heated continuously. A small test flame was introduced into the cup at regular intervals, and the flash point was taken as the lowest temperature at which application of the test flame causes the vapour above the sample to ignite with a distinct flash inside the cup.

#### 2.3.2 Gross calorific value

The gross heat of combustion of a fuel at constant volume is the number of heat units measured as being liberated at 25°C when unit mass of the fuel is burned in oxygen saturated with water vapour in a bomb under standard conditions(Indian Standard Methods of Test for Petroleum and its Products (P6)). A bomb calorimeter was used for this experiment. A weighed quantity of the biofuel sample was burned in oxygen inside a bomb calorimeter under controlled conditions. The gross calorific value was calculated from the mass of the sample and the rise in temperature.

#### 2.3.3 Cloud point

:It is the temperature at which a cloud of wax crystals first appears in a liquid when it is cooled under specified conditions (Indian Standard Methods of Test for Petroleum and its Products (P10)) A sample of biofuel was cooled slowly and examined periodically. The temperature at which a cloud was first observed at the bottom of the test jar was recorded as the cloud point.

## 3. RESULTS AND DISCUSSIONS

### 3.1 Extraction, transesterification and FAME analysis

Figure 1 a and b below show the Fame analysis of the roots and stems of phumdis respectively.

**Fig. 1.**
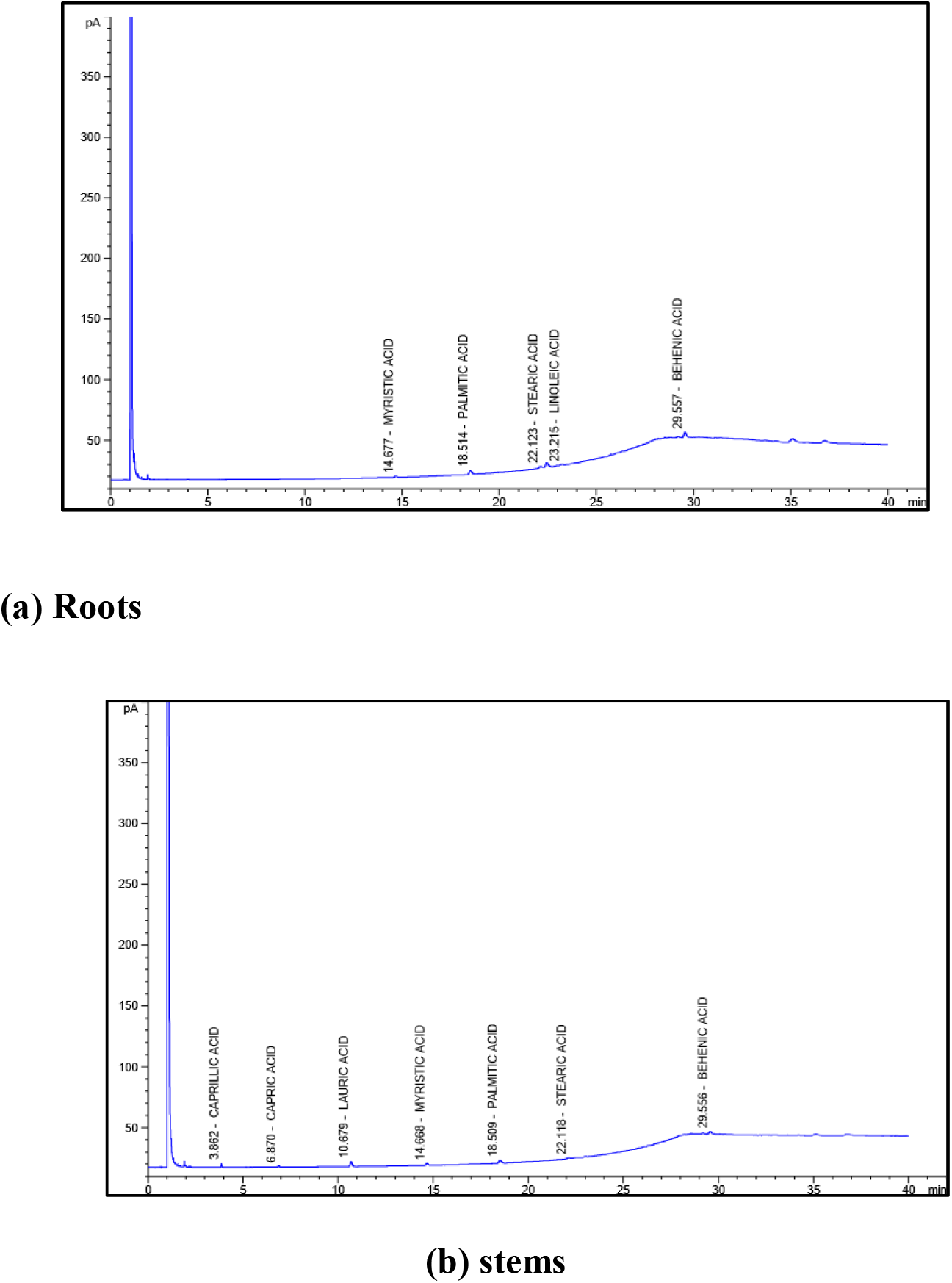
**FAME analysis of the** roots and stems of Phumdis. The results indicate the presence of a high amount of behenic acid, stearic acid and palmitic acid in both the samples.

#### 3.3.1 Flash point

This is an important property which determines flammability of the material. It should also be taken into account during transporting, handling and storage. The flash point of the sample was found to be below 5 °C which indicates that it is a very inflammable liquid. However its safety can be increased by blending it with diesel at various proportions.

#### 3.3.2 Gross calorific value

It is the total energy released in the form of heat when the sample undergoes complete combustion in the presence of oxygen hence making it an important property in the determination of fuel. The gross calorific value of the sample was found to be 4548 cal/g or 19.029 kJ/kg.

#### 3.3.3 Cloud point

With the presence of waxes,the oil thickens and clogs the engine. It is a serious threat issue in cold countries. The cloud point of the sample was found to be 11°C.

## 4. CONCLUSION

As we have seen, the high calorific value, cloud point, flashpoint of phumdis indicate that it can be used as an alternative biofuel. The use of phumdis promises to be a win – win situation by helping in maintaining the lake ecosystem as well as generating biofuel. It is an innovative use of a waste land that burdens the eastern part of India. It is a way forward in the field of renewable energy encouraging more research in types of biomass that have the potential to produce energy.

## DECLARATIONS

### ETHICS APPROVAL AND CONSENT TO PARTICIPATE

Not applicable

### CONSENT FOR PUBLICATION

Not applicable

### AVAILABILITY OF DATA AND MATERIALS

Not applicable

### AUTHORS’ CONTRIBUTIONS

SK, AS and JJ are responsible for pre-treatment of phumdis, culturing the biomass and extracting the biofuel from it. They also performed analytical tests of the biomass (Phumdis) to determine its potential for producing biofuel.

DK and KT are responsible for mechanical testing of fuel produced flash point, cloud point and gross-calorific value.

SV and CKCB are primary supervisors who provided scientific knowledge in bioprocessing and mechanical testing respectively to successfully complete the project.

## FUNDING AND ACKNOWLEDGEMENTS

Financial support from TEQIP –II, BMSCE, Bangalore for this research is gratefully acknowledged.

